# Microampere electric currents caused bacterial membrane damage and two-way leakage in short time

**DOI:** 10.1101/2020.03.13.991067

**Authors:** V R Krishnamurthi, A Rogers, J Peifer, I Niyonshuti, J Chen, Y Wang

## Abstract

Physical agents such as low electric voltages and currents have recently gained attention for antimicrobial treatment due to their bactericidal capability. Although microampere electric currents were shown to suppress the growth of bacteria, it remains unclear to what extent the microampere currents damage bacterial membrane. Here, we investigated the membrane damage and two-way leakage caused by microampere electric currents (≤ 100 μA) in a short time (30 min). Based on MitoTracker staining, propidium iodide staining, filtration assays, and quantitative single-molecule localization microscopy, we found that microampere electric currents caused significant membrane damages and allowed two-way leakages of ions, small molecules and proteins. This study paves the way to new development and antibiotic applications of ultra-low electric voltages and currents.

**Statement of Significance:** Previous studies showed that treating bacteria with milliampere electric currents for 72 hours led to significant damages of the bacterial membrane. However, it remains unclear to what extent membrane damages and two-way (i.e. inward and outward) leakages are caused by lower electric currents in a shorter time. In this work, we set out to answer this question. We carried out several assays on the bacteria treated by microampere electric currents of ≤ 100 μA for 30 min, including MitoTracker staining, propidium iodide staining, filtration assays, and quantitative single-molecule localization microscopy. We found and quantified that the membrane damages were caused by microampere electric currents in half an hour and allowed two-way leakages of ions, small molecules, and proteins.

## Introduction

As antibiotic resistance of bacteria has become one of the biggest threats to public health (1), alternatives to antibiotics have been attracting broad interest and attention (2, 3). Physical methods, such as sunlight/UV, fire, drying, high temperature, and high pressure, have played important roles in sterilization and disinfection from very early times of human history (4). Since the 1960s, electric voltages, currents, and fields have been explored as physical means for suppressing the growth of, and/or killing, bacteria. While most of the early studies focused on the bactericidal effects of high electric voltages and currents (5–8), it was found, more recently, that low electric voltages and currents can also effectively kill bacteria and biofilms (9–27). For example, Pareillexu and Sicard investigated the effects of low electric currents ranging from 10 to 200 milliampere (mA) on the viability *Escherichia coli* (*E. coli*) bacteria and found that currents as low as 25 mA could kill the bacteria (27). The lethal activity of mA currents were confirmed by various later reports (9, 17–19, 25, 26). In addition, many studies showed that ultra-low electric current at microampere (μA) were bactericidal (10, 13, 18, 20, 21, 24); even electric field (without current) can inhibit the growth of planktonic bacteria (16). Furthermore, electric currents at μA and/or mA have been applied to, and shown to be effective for treating biofilms (11, 12, 14, 15, 18, 22–24, 28).

Efforts have been made toward understanding the antimicrobial mechanisms of low electric voltages and currents. Possible mechanisms include membrane damage and disruption, reduction in ATP production and enzymatic activities (17, 26), and generation of reactive oxygen species (ROS) (13–15). Membrane damages caused by high electric voltages have been well known for a long time. For example, electroporation is a commonly used microbiological technique to deliver DNA and proteins into bacteria and cells by applying high electric voltages at kilovolts (kV) (29–31). More recently, transmission electron microscopy showed that treating bacteria at low electric currents of 5 mA for ≥ 72 hours led to significant damages, based on which leakage of cellular contents and influx of toxic substances through the damaged membrane were suggested (17). An interesting follow-up question is to what extent membrane damages and twoway (i.e. inward and outward) leakages are caused by microampere electric currents in a shorter time. Although previous results showed that microampere electric currents were effective for suppressing the growth of bacteria, it remains unclear to what extent the microampere currents damage bacterial membrane.

In this work, we set out to answer this question by investigating possible membrane damages of bacteria after subjecting the bacteria to microampere electric currents of ≤ 100 μA (corresponding to voltages ≤ 2.5 V) for 30 min. From MitoTracker staining experiments, we observed that treated bacteria showed much higher fluorescent intensities, indicating that bacterial membrane was affected by the electric currents as low as 50 μA (or ≈ 1.5 V). In addition, using propidium iodide (PI) staining and filtration assays, we found that the membrane damages allowed two-way leakages of ions, small molecules, and even proteins. This observation was confirmed by quantitative single-molecule localization microscopy, which allowed us to count the number of histone-like nucleoid structuring (H-NS) proteins inside individual bacteria, confirming that the number of H-NS proteins per bacterium decreased as the applied microampere electric currents increased. Further quantitative analysis suggested that the organization and clustering of the H-NS proteins were affected by the applied electric currents and voltages. This study highlights that microampere electric currents (≤ 100 μA) in short time scales (e.g. 30 min) caused significant membrane damages to allow two-way leakages of ions, small molecules and proteins through the bacterial membrane.

## Materials and Methods

### Bacterial strain and growth

Two K-12 derived *E. coli* strains were used in this study: the first strain is MG1655 (from the Yale Coli Genetic Stock Center) (32, 33), and the second strain has the *hns* gene on the bacterial DNA fused to *meos* gene. The second strain expresses fluorescent H-NS-mEos3.2 fusion proteins (34), facilitating the fluorescence-based filtration assay and the quantitative single-molecule localization microscopy (34–38).

The bacteria were grown at 37°C overnight in defined M9 minimal medium, supplemented with 1% glucose, 0.1% casamino acids, 0.01% thiamine and appropriate antibiotics with orbital rotation at 250 rpm (35–37, 39). On the second day, the overnight culture was diluted by 50 to 100 times into fresh medium so that the OD_600_ was 0.05 (35–37, 39). The fresh cultures were grown again at 37°C in culture tubes equipped with two sterile aluminum electrodes with orbital rotation at 250 rpm without electric voltages or currents (i.e., the wires were shorted, Fig.1A). The resistance of the bacterial cultures (≈ 32 kΩ) was measured using a multimeter. When the fresh bacterial culture reached OD_600_ ≈ 0.3, low DC voltages were applied to the culture for 30 min (0V – untreated negative control, 0.5, 1, 1.5, 2, and 2.5 V, Fig. 1B). The corresponding currents were 0, 16, 31, 47, 63, and 78 μA, respectively. After the 30 min treatment, the bacteria were used for the experiments and quantifications as described below.

**Figure 1.**
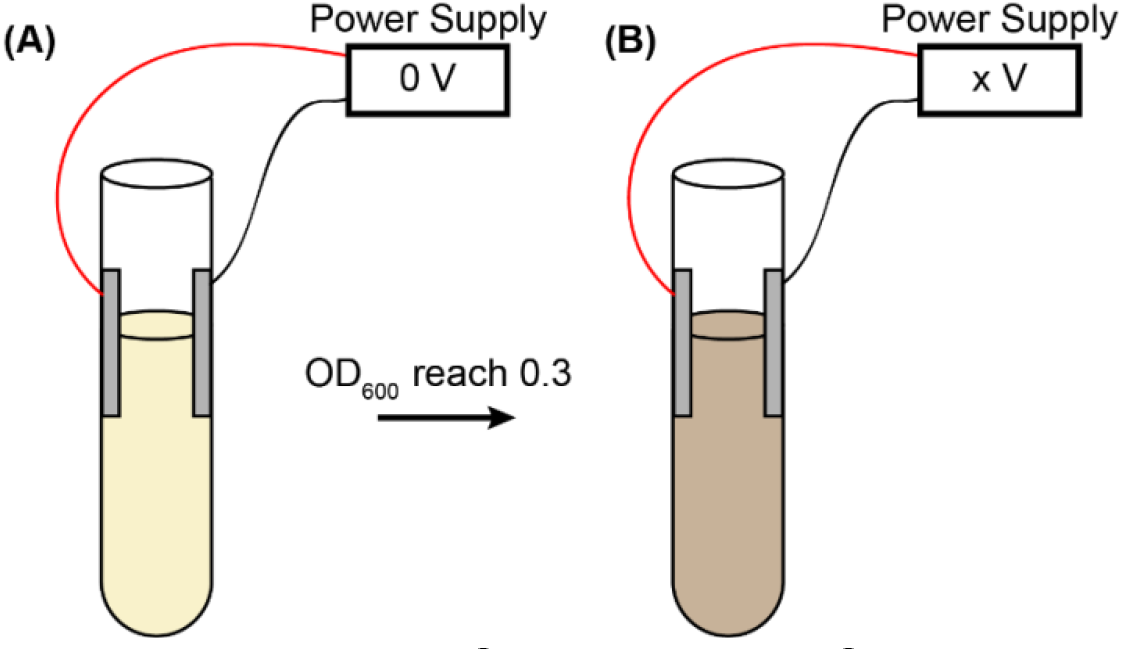
Treating bacteria with low DC voltages. (A) Overnight bacterial culture was diluted in fresh medium and regrew until OD600 reached 0.3 without applying DC voltage (0 V). (B) Applying low DC voltage (0.5, 1.0, 1.5, 2.0, and 2.5 V) on the bacterial culture in culture tubes for 30 min, followed by quantifications.

### MitoTracker staining and quantification

To stain the membrane of bacteria, MitoTracker Green FM dyes (Thermo Fisher Scientific) were added to the untreated and treated bacteria at a final concentration of 300 nM, and incubated in a shaking incubator (250 rpm, 37°C) for 30 min (40). 8 μL of the stained bacteria were transferred to 5mm × 5mm agarose pads (3% in 1X PBS, ~1 mm thick) for mounting. The agarose pads with bacteria were flipped and attached to clean coverslips (cleaned with sonication in 1 M NaOH, 100% ethanol, and ultra-pure water sequentially). Chambers were then constructed by sandwiching rubber o-rings between the coverslips and cleaned microscope slides. The chambers were sealed using epoxy glue and then mounted on a microscope for fluorescence imaging (excitation = 488 nm). From the acquired images, the average fluorescence intensities of 100 bacteria treated at each voltage were measured using ImageJ (41,42).

### Propidium iodide (PI) staining and quantification

To measure possible inward leakage due to membrane damage, the untreated and treated bacteria were first fixed by 3.7% formaldehyde (Sigma-Aldrich) and stained with PI (G-Biosciences) (43, 44). The PI-stained bacteria were mounted on agarose pads and prepared for fluorescence imaging similar to the MitoTracker experiments, except that the excitation wavelength was changed to 532 nm. The stained cells were counted and their percentages were calculated based on the acquired fluorescence images.

### Filtration assays

Filtration assays to assess the leakage of cellular contents due to membrane damages were performed similar to Ref. (45). Briefly, the untreated and treated bacteria (expressing H-NS-mEos3.2 fusion proteins) were filtered by 0.2 μm filters (VWR International LLC), resulting in filtrates constituting leaked cellular contents due to membrane damage, in addition to proteins and molecules from the culture media. For each sample, the filtrate was aliquoted into 96-well plate (200 μL per well), followed by measuring the absorbance at 280 nm and fluorescence at 525 nm (excitation = 498 nm) on a multi-mode microplate reader (Synergy H1, BioTek).

### Quantitative single-molecule localization microscopic assay

Quantitative single-molecule localization microscopy (46) on the bacteria subjected to microampere electric currents was performed following our previous work (35–37, 39). Briefly, the untreated and treated bacteria were first fixed by 3.7% formaldehyde for 30 min at room temperature and harvested by centrifugation (1000 g for 10 min). The harvested cells were resuspended in 1X phosphate-buffered saline (PBS) buffer. The centrifugation and resuspension were repeated for three times. The prepared bacteria were mounted on agarose pads for imaging. The single-molecule localization microscope was home-built on an Olympus IX-73 inverted microscope with an Olympus TIRF 100X N.A.=1.49 oil immersion objective. The microscope and data acquisition were controlled by Micro-Manager (47, 48). A 405 nm laser and a 532 nm laser from a multi-laser system (iChrome MLE, TOPTICA Photonics) were used to “activate” and excite the H-NS-mEos3.2 fusion proteins in bacteria. Emissions from the fluorescent proteins were collected by the objective and imaged on an EMCCD camera (Andor) with an exposure time of 30 ms (the actual interval between frames was 45 ms). The effective pixel size of acquired images was 160 nm, while the field of view was 256 × 256 pixels.

The resulted movies (20,000 frames) were analyzed with RapidStorm (49), generating x/y positions, x/y widths, intensity, and background for each detected fluorescent spot. Spots that are too dim, too wide, or too narrow, were rejected (50), followed by drift-correction using a mean cross-correlation algorithm (51). The localizations that appear in adjacent frames and within 10 nm to each other were regrouped as a single molecule (52). The resultant localizations were used for reconstruction of super-resolved images and for further quantitative analysis.

The localizations of H-NS molecules from each acquisition were first manually segmented into individual bacterial cells using custom MATLAB programs (35). Analysis and quantification of the spatial organization of H-NS proteins were performed on individual cells. Following the segmentation of cells, the number of localizations per bacterium *N_p_* was counted (35). In addition, Voronoi-based quantification and clustering analysis were performed using custom MATLAB programs following Levet et al. (53), from which we estimated the molecular density of H-NS proteins *ρ*, the mean inter-neighbor distances *Δ_ave_*, the number of clusters per bacterium *N_cl_*, the number of proteins per cluster *N_pc_*, the area of clusters of H-NS proteins *A_cl_*, and the fraction of proteins forming clusters *φ_cl_* (35, 53).

## Results

### Membrane damage caused by microampere electric currents

Microampere electric currents were applied to the bacterial culture by applying low electric DC voltages. As the resistance (R) of the bacterial culture was measured to ≈ 32 kΩ, DC voltages (V) in the range of 0 – 2.5 V resulted in currents (I) of 0 – 78 μA (I = V / R). After treatment for 30 min, we stained the bacterial membrane by MitoTracker Green FM dyes (40). As shown in Figs. 2A and 2B, the intensity of the bacterial membrane became higher after subjecting the bacteria to the microampere electric currents. To quantify this observation, we estimated the mean intensities of the bacteria and examined the dependence of the mean intensity on the voltage (Fig. 2C). The intensity increased slightly at 0.5 V or 16 μA, indicating changes in the bacterial membrane. With ≥ 1.5 V or 47 μA, the average intensity increased to ≥ 4-fold, while many bacterial showed extremely high brightness (up to 19 times brighter) compared to the negative control, suggesting that many of the treated bacteria had significant changes in their membrane. As previous studies showed that, although controversy exists (54, 55), the fluorescence of membrane-incorporated MitoTracker Green dyes depends on the membrane potential (56, 57), it is possible that the microampere electric currents altered the bacterial membrane potential.

**Figure 2.**
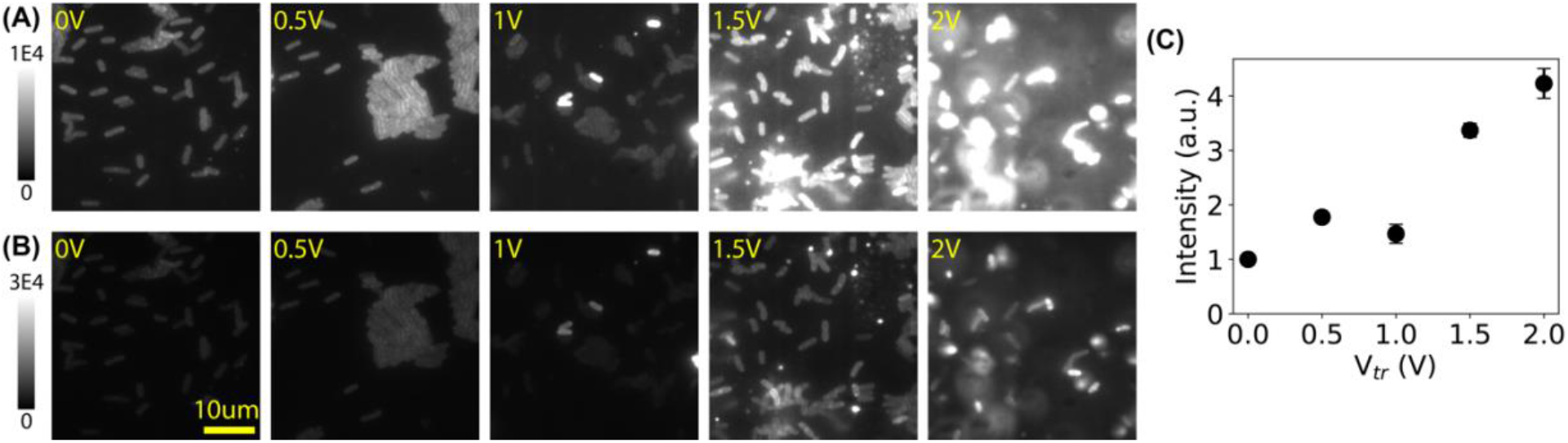
Membrane damage in bacteria caused by microampere electric currents. (A) Representative fluorescence images of bacterial membrane stained by MitoTracker Green FM dyes, after treatment of low DC voltages at 0 V (untreated negative control) to 2.0 V (corresponding to 0 – 63 μA). (B) The same images from panel A, but with a larger intensity scale. (C) Dependence of the mean fluorescence intensity (rescaled by the negative control) on the applied voltage.

### Inward leakage due to membrane damage caused by microampere electric currents

It was previously proposed that membrane damages caused by low electric currents allowed influx of toxicants (17). To test this possibility with microampere electric currents, propidium iodide (PI) (43, 44), with a size of ≈ 1 nm, was used to stain the DNA of the bacteria after treatment by microampere electric currents. The rationale is that, if the electrically induced membrane damages allow PI to enter the bacteria to stain DNA, influx of ions and other small organic molecules would also be possible. Representative images of untreated (0 V or 0 μA) and treated (0.5 – 2.5 V or 16 – 78 μA) bacteria are shown in Fig. 3A, where the fluorescent images from PI staining (red) are superposed on the inverted phase contrast images (blue). It was observed that more bacteria were stained by PI at higher voltages and currents. Quantifying the percentage of PI-stained bacteria (P_pi_) showed that the percentage of membrane-damaged bacteria increased quickly from 20% to ≈ 80% at 1.5 – 2 V or 47 – 63 μA (Fig. 3B). This observation confirmed that the membrane damages caused by microampere electric currents are significant enough to enable inward leakage of at least ions and small molecules with a size of ≤ 1 nm.

**Figure 3.**
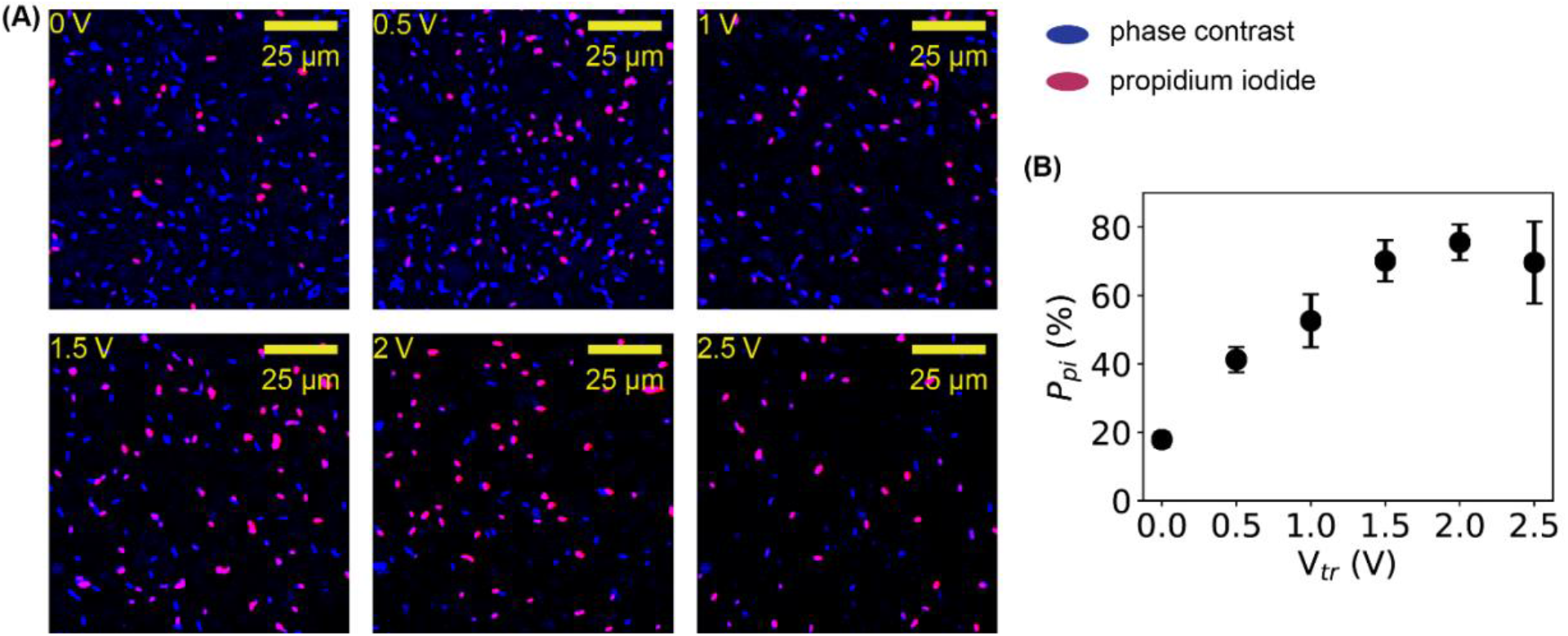
Quantification of inward leakage due to membrane damage by PI staining. (**A**) Representative fluorescence images (red) of bacteria stained by PI before and after treatment of electric treatment, superposed on the inverted phase contrast images (blue) of the bacteria in the same fields of views. (**B**) Dependence of the percentage of PI-stainable bacteria on the applied voltage.

### Outward leakage due to membrane damage caused by microampere electric currents

Our PI-staining result showed that influx of ions and small organic molecules (≤ 1 nm) was possible due to membrane damages caused by microampere electric currents, which also implies that small cellular contents, including amino acids, could leak out of the bacteria. Another question is whether biological macromolecules, such as proteins and nucleic acids, leak out of the bacteria due to the membrane damages caused by microampere electric currents. To answer this question, we adopted a filtration assay that was used to study the membrane damages caused by carbon nanotubes (45). Briefly, the treated bacteria were filtered by 0.2 μm filters, and the absorbance of the filtrates was measured at 280 nm, at which both nucleic acids and proteins absorb. We then estimated the relative changes by 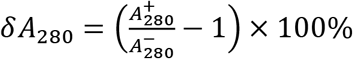, where 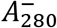 and 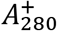 are the absorbances at 280 nm of the filtrates for bacteria before and after microampere current treatment, respectively. A slight increase (~1%) was observed (Fig. 4A), providing weak evidence of possible outward leakage of macromolecules. Because the “background” from the culture media could contribute significantly to the absorbance, it is likely that the measured increase in the absorbance was underestimated.

**Figure 4.**
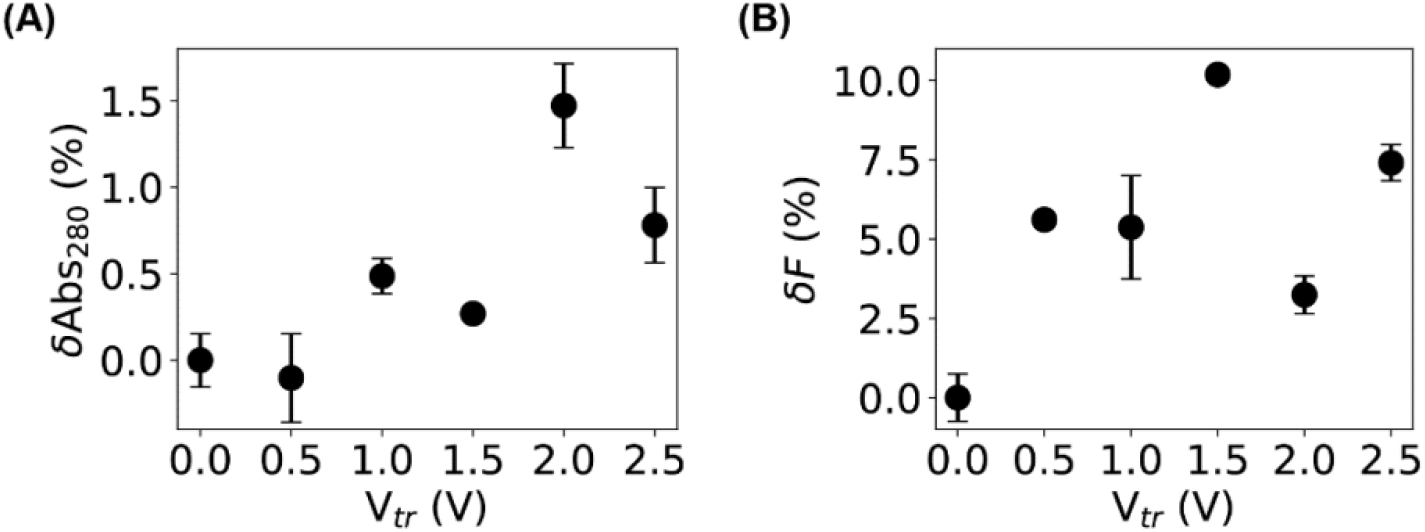
Quantification of outward leakage of bacterial content due to membrane damage by the filtration assays. (**A**, **B**) Relative changes in the (**A**) absorbance at 280 nm and (**B**) fluorescence of the filtrates of bacteria samples before and after electric treatment.

To lower the “background” due to the culture media, a modified filtration assay based on fluorescence was performed using an *E. coli* strain that expresses histone-like nucleoid structuring (H-NS) proteins fused to mEos3.2 fluorescent proteins (34). The fluorescence-based filtration assay allowed us to measure the outward leakage of proteins more specifically. By quantifying the relative changes in the fluorescence intensities of the filtrates, 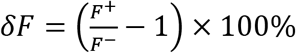, where *F^−^* and *F^+^* are the fluorescence of the filtrates for samples before and after microampere current treatment, respectively, we observed an increase of ~10% for bacteria treated at 2.5 V or 78 μA. This observation suggested that the membrane damages caused by the microampere electric currents were significant enough to allow proteins to leak out of the bacteria.

### Quantification of protein leakage by single-molecule localization microscopy

The filtration assays suggested that proteins leaked out of bacteria after treating the bacteria by ≤ 100 μA DC currents. To further confirm such protein leakage, we measured the number of the histone-like nucleoid structuring (H-NS) proteins inside individual bacteria before and after microampere current treatment for 30 min using single-molecule localization microscopy (SMLM) (46, 50, 58, 59). SMLM is one type of super-resolution fluorescence microscopy that localizes individual molecules of interest with a precision of ≤10 nm (60); therefore, it not only produces super-resolved images with high spatial resolution, but also provides a convenient way to count the number of molecules. Without applying sophisticated algorithms (61–63), the number of molecules of interest is on average proportional to the intensity in the super-resolved images or the number of localizations obtained by SMLM (39).

We performed single-molecule localization microscopy on untreated and treated bacteria, using the *E. coli* strain expressing H-NS-mEos3.2 fusion proteins, as mEos3.2 fluorescent proteins are photoactivable and allow super-resolution imaging (34, 35). Representative images of H-NS proteins in untreated and treated bacteria are shown in Fig. 5A. For the untreated negative control, H-NS proteins were organized as clusters inside the bacteria (first column in Fig. 5A and 5B), consistent with previously reported results (35, 36, 38). After subjecting the bacteria to DC voltages of 1 – 2.5 V (or currents of 31 – 78 μA), the intensities on the super-resolved images became significantly dimmer (columns 3–6 in Fig. 5A and 5B). As the intensity in super-resolved images correlates with the probability of localizing the molecules of interest, this result indicates that less H-NS proteins were present in the bacteria after treating the bacteria with microampere electric currents.

**Figure 5.**
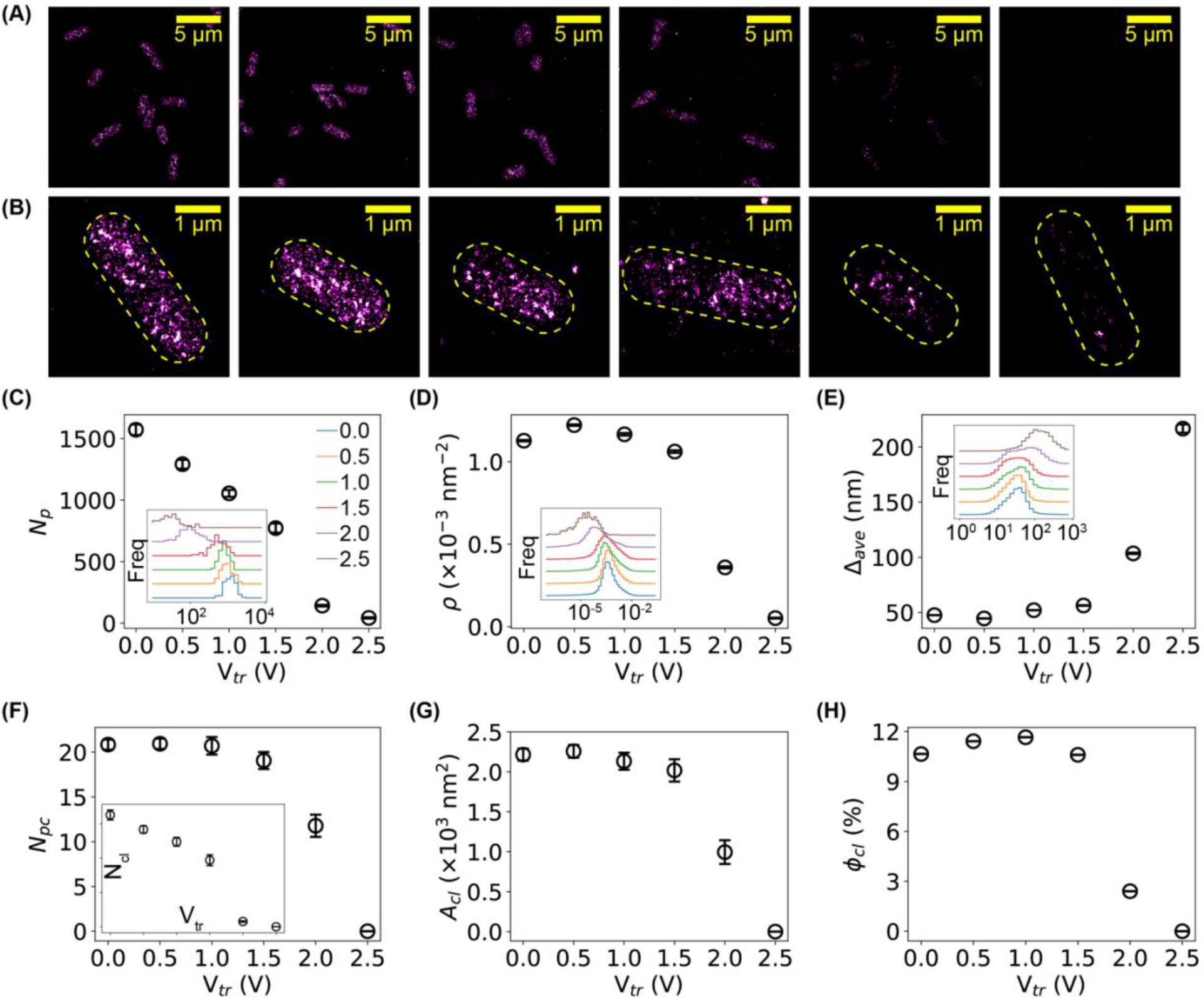
Quantification of outward leakage of bacterial content due to membrane damage by single-molecule localization microscopy. (**A**) Representative super-resolved images of H-NS proteins in *E. coli* bacteria before and after electric treatment. Scale bar = 5 μm. (**B**) Zoom-in view of super-resolved images showing individual bacteria. Scale bar = 1 μm. (**C – E**) Dependence of (C) the number of detected H-NS proteins per cell Np (D) the molecular density ρ, and (E) the average inter-molecular distance *Δ_ave_* on the applied voltage. Insets are the corresponding distributions of the three quantities. Error bars stand for the standard error of the mean (SEM). (**F – H**) Dependence of (F) the number of clusters per cell N_cl_ (inset) and the number of proteins per cluster N_pc_, (G) the area of clusters A_cl_, and (H) the fraction of clustering proteins per cell *ϕ_cl_* on the applied voltage. Error bars stand for SEM.

To quantify this result, we segmented the bacteria (35) and counted the number of localizations of H-NS proteins in each cell, *N_p_*. Due to the stochastic nature in the activation and detection of the mEos3.2 fluorescent proteins, *N_p_* is expected to be proportional to, on average, the number of H-NS proteins in the bacteria (35, 39). The histograms of *N_p_* are shown in the inset of Fig. 5C, where the baselines of the histograms were vertically shifted for better visualization of the differences. It is clear that the peaks of the *N_p_* distributions translated to the left as the applied voltage increased, suggesting that the number of H-NS proteins per bacterial cell decreased when subjecting the bacteria to the low DC voltages and currents. We also calculated the mean of *N_p_* and standard error of the mean (SEM) for each sample. Note that, as the distributions of *N_p_* were bell-shaped in the linear-log scale (inset of Fig. 5C), we estimated the mean of *log(N_p_)* and then calculated the mean of *N_p_* from *N_p_* = *exp*(*log(N_p_)*). As shown in (Fig. 5C), the average number of H-NS proteins per cell decreased nearly linearly (Fig. 5C). It is also noted that the observed decrease in the number of H-NS proteins per bacterial cell is unlikely due to cellular responses (e.g., lower expression level of H-NS proteins) for several reasons. First, the treatment time (30 min) is relatively short, comparing to the degradation time of proteins in bacteria (hours) (64). Second, our previous work on H-NS proteins in bacteria treated by antibiotic silver ions and silver nanoparticles showed that the number of H-NS proteins did not decrease within several hours (35). Therefore, the microscopic data on individual bacteria directly suggested that H-NS proteins leaked out of the bacteria after microampere electric current treatment.

We examined how the spatial organization of the H-NS proteins was affected by the electric treatment based on Voronoi diagram tessellation (35, 53). We computed the molecular density (*ρ*) of the H-NS proteins and found that the distribution of *ρ* shifted to the left, indicating the density decreased as the applied voltage increased (inset of Fig. 5D), which is expected as a result of the decrease of the number of H-NS proteins per cell. Interestingly, the mean of the molecular density started to decrease at 1.5 – 2 V (or 47 – 63 μA) as shown in Fig. 5D, which is different from the dependence of the number of H-NS proteins per cell on the applied electric voltages and currents (Fig. 5C). We also observed a shift to the right in the distribution of the mean inter-neighbor distances *Δ_ave_* (inset of Fig. 5E), suggesting that the inter-molecular distances for some H-NS proteins became larger when bacteria were treated by low DC voltages and currents, consistent to the result of the molecular density. In addition, we identified the clusters of H-NS proteins based on Voronoi diagrams (35, 53) and counted the number of clusters per bacterial cell (*N_cl_*) and the number of proteins per cluster (*N_pc_*). We found that both *N_cl_* and *N_pc_* decreased after treating the bacteria with DC voltages and currents (Fig. 5F). Furthermore, by quantifying the area of clusters of H-NS proteins, we observed that the H-NS clusters became smaller (Fig. 5G). Lastly, we examined the fraction of H-NS proteins forming clusters in individual bacteria, *ϕ_c_* = *N_cp_*/*N_p_*, where *N_p_* is the number of localizations (i.e., detected proteins) per cell, and *N_cp_* the number of proteins that form clusters in the cell. We observed that the clustering fraction decreased from 10% for the negative control to 0% at 2.5 V or 78 μA (Fig. 5H).

### Conclusions and Discussions

To summarize, we investigated the membrane damage of bacteria and the two-way leakage caused by microampere electric currents. We observed that bacteria subjected to ≥ 47 μA currents for 30 min showed much higher intensities with MitoTracker staining, suggesting that the bacterial membrane was altered by the microampere electric currents. The microampere current caused membrane damages were large enough to allow PI molecules to enter the bacteria, suggesting that inward leakages of ions and small molecules were possible. In addition, based on filtration assays and super-resolution imaging results, we found that the membrane damages were so significant that proteins leaked out of the bacteria. More importantly, using histone-like nucleoid structuring (HNS) proteins as an example, we quantified the decrease in the number of H-NS proteins per bacterial cell and characterized the changes in the spatial organization of the H-NS proteins caused by the electric treatment. This study highlights that treating bacteria with electric currents at ≤ 100 μA for 30 min caused significant membrane damage and led to two-way leakages of ions, small molecules and proteins.

It was noted previously that the bactericidal effects of electric voltages and currents are complex, and involve various interactions between the bacteria, electricity, electrode materials and medium (17). For example, in addition to the electrical effects, metal ions released from the electrodes into the medium are likely to affect the growth of bacteria, which has been shown in the literature (65). In addition, ROS generated by the electric voltages and currents could be another significant contributor (13–15). The current study did not aim to distinguish the contributions of these different effects, while these are interesting questions worth pursuing in future investigations. On the other hand, we argue that heating by the electric voltage/current is unlikely to cause damages to the bacteria in this study, because the used electric power is very low. Considering the resistance of the bacterial culture was ≈ 32 kΩ, the electric power was below 200 μW; at this power, it would take more than one day to heat up the bacterial culture (5 mL) by one degree.

It is worthwhile to highlight that the electric power leading to serious membrane damages of bacteria is very low, which is expected to facilitate the use of microampere electric currents (and low electric voltages) for antibiotic applications. For example, commonly used household batteries can provide the needed voltages to kill bacteria. More importantly, solar panels have an output power of ~10 mW per cm_2_ (66); therefore, a solar panel of 1 cm_2_ can easily generate the needed electric power for damaging bacterial membranes, suppressing the growth of bacteria and/or killing bacteria. We anticipate that this study intrigues new development and antibiotic applications of low electric currents.

## Acknowledgment

We thank Dr. Joshua N. Milstein for the generous gift of the *E. coli* strain expressing mEos3.2-HNS fusion proteins. This work was supported by the University of Arkansas, the Arkansas Biosciences Institute (Grant No. ABI-0189, No. ABI-0226, No. ABI-0277), and the National Science Foundation (Grant No. 1826642). A.R. and J.P. were supported through the Research Experience for Undergraduate program funded by the National Science Foundation (Grant No. 1460754). We are also grateful for supports from the Arkansas High Performance Computing Center (AHPCC), which is funded in part by the National Science Foundation (Grants No. 0722625, 0959124, 0963249, 0918970) and the Arkansas Science and Technology Authority.

## Author Contributions

Y.W. and J.C. conceived and planned the experiments. V.R.K. performed all the experiments. V.R.K. J.P and A.R. performed the imaging. V.R.K and Y.W. performed data analysis and interpretation. I.I.N characterized solutions. Y.W., and J.C supervised the project.

